# Entering and Exiting Awareness are Asymmetrical: Evidence from tracking Continuous Flash Suppression

**DOI:** 10.64898/2026.10.18.759988

**Authors:** Cameron K. Phan, Reuben Rideaux, David Alais

## Abstract

This study provides a model for obtaining cleaner threshold estimates of entry into and exit from conscious awareness measured with the popular breaking continuous flash suppression (bCFS) and newer tracking CFS (tCFS) paradigms. In both, a suppressed target increases in contrast until it breaks into awareness. tCFS also measures exit thresholds by reversing the contrast change. Both CFS paradigms require a response when the threshold is reached and thus involve a reaction-time component. During this period, target contrast continues to change and threshold estimates are inflated (the “overshoot” problem). Here, we measure reaction-time components for both increasing (entering awareness) and decreasing targets (exiting awareness). We find overshoot is asymmetrical, being greater for stimuli entering (breakthrough thresholds) than for exiting awareness (suppression thresholds). To solve “the overshoot problem”, we introduce a mathematical model that quantifies reaction-time components so they can be partialled out to produce clean breakthrough and suppression thresholds. The model can also be applied to previously published CFS studies. The model works for simple target detection and two-alternative target discrimination, even in the continuous tracking paradigm of tCFS. With the timing corrections applied, changes in pupil diameter around transitions in awareness are shown to align with the corrected time points, most clearly when targets exit awareness. Even after correction, the contrast decrease required for a target to exit awareness was greater than the increase required for it to enter awareness.

## Introduction

Among the totality of things encoded in the brain, there are some that we are aware of and much that we are not. Our “phenomenal consciousness” (Block, 1995) or “conscious content” (Seth, 2009) has passed through a kind of bottleneck limiting access to consciousness and we are aware of only a small fraction of neural processing (Dehaene & Naccache, 2001; Sklar et al., 2021). In studies of visual consciousness, an enduring quest has been to discover how much visual processing can occur outside of conscious awareness and what kinds of images are prioritized to gain access to conscious expression. Although a number of methods can be employed to study visual suppression and access to awareness (Kim & Blake, 2005), binocular rivalry (Alais & Blake, 2005, 2015) and continuous flash suppression (CFS: (Tsuchiya & Koch, 2005)) have been the most widely used. Binocular rivalry has a long history (Alais et al., 2000; Breese, 1909; Wheatstone, 1838) and was touted as a key method for understanding visual consciousness in the 1990s (Baker, 2010; Crick, 1996; Logothetis, 1998).

More recently, however, CFS has gained in popularity as a way to study visual processing outside of awareness and is now the dominant approach (Pournaghdali & Schwartz, 2020; E. Yang et al., 2014). Both binocular rivalry and CFS are forms of interocular suppression caused by the eyes viewing conflicting stimuli that prevent binocular fusion. This conflict triggers interocular suppression so that only one eye’s view is perceived at a time and the other is suppressed. In binocular rivalry, this monocular suppression will typically last a second or so before a switch occurs and the other eye becomes suppressed, releasing the formerly suppressed eye from suppression. The chief advantage of CFS is that rather than using two static images it presents a rapidly flickering pattern to one eye (the masker or ‘Mondrian’) which suppresses a static and low-contrast image (the ‘target’) presented to the other, a combination that produces strong and long-lasting suppression (Tsuchiya et al., 2006; Tsuchiya & Koch, 2005).

CFS is now much more widely used than binocular rivalry, due mainly to a very popular variant known as ‘breaking-CFS’ (bCFS: (Jiang et al., 2007; Stein, Hebart, et al., 2011). Unlike classical CFS, in which a suppressed target can remain out of awareness for long periods (several tens of seconds, even a minute or so), suppression in bCFS is much shorter because the target is initially presented at low contrast but quickly ramps up in contrast until it is strong enough to break suppression and enter awareness (a matter of a few seconds). The time required for the target to reach breakthrough is the dependent variable. Often different target images are compared and a faster breakthrough time for one image relative to another is interpreted as that image having priority access to awareness or undergoing greater processing outside of awareness before entering awareness (Costello et al., 2009; Jiang et al., 2007; Mudrik et al., 2011; Stein, Senju, et al., 2011; E. Yang et al., 2007; Y.-H. Yang & Yeh, 2011).

One limitation of the bCFS approach is that some images are simply more salient to neurons in early visual cortex than others, due to factors such as image contrast and spatial structure (Bex et al., 2009; Meese et al., 2017; Peli, 1990). For this reason alone, some images would be expected to break through sooner than others by eliciting a greater neural response. A recent variation known as ‘tracking-CFS’ (tCFS: (Alais et al., 2024; Alais & Kim, 2025) deals with this by finding each image’s own suppression threshold. This is done by ramping down an image’s contrast from a high (and visible) starting point until it succumbs to suppression from the Mondrian in the other eye. In effect, tCFS aligns images by their suppression threshold to take account of salience variations among images and focuses on the contrast increment required to reach breakthrough. Since all tCFS measurements are made on a log-contrast scale, the increment is actually a ratio in which breakthrough is x-times greater than suppression threshold. Breakthrough thresholds for different images can then be compared equitably by ramping contrast up from their suppression threshold and calculating the proportionate change in contrast required to break into awareness (Alais et al., 2024).

Part of the appeal of bCFS and tCFS is that they are simple to implement and give quick and efficient measures of breakthrough. tCFS has an advantage in also providing a measure of suppression threshold, thus allowing easy calculation of how deeply an image is suppressed from awareness by comparing the difference between breakthrough and suppression thresholds (expressed in terms of log contrast). This informative variable is known as ‘suppression depth’ and was often measured in binocular rivalry studies (Alais & Melcher, 2007; Blake & Camisa, 1979; Lunghi & Alais, 2015; Makous & Sanders, 1978; Nguyen et al., 2003) but very rarely in CFS (the two exceptions among hundreds of papers are (Tsuchiya et al., 2006; E. Yang & Blake, 2012)). Both CFS paradigms, however, suffer from a limitation related to using images undergoing contrast ramping. Specifically, once the observer notices the target has broken into awareness (or disappeared into suppression, in the case of tCFS), there is a reaction-time period in which the contrast ramp continues while the observer plans and executes a motor response to report the change in image visibility. Consequently, there is an “overshoot problem” that applies to both bCFS and tCFS that causes the breakthrough threshold to be overestimated. In addition, overshoot in the tCFS paradigm will produce underestimated suppression thresholds (**Figure 1a**), with a consequent inflation of suppression depth.

**Figure 1.**
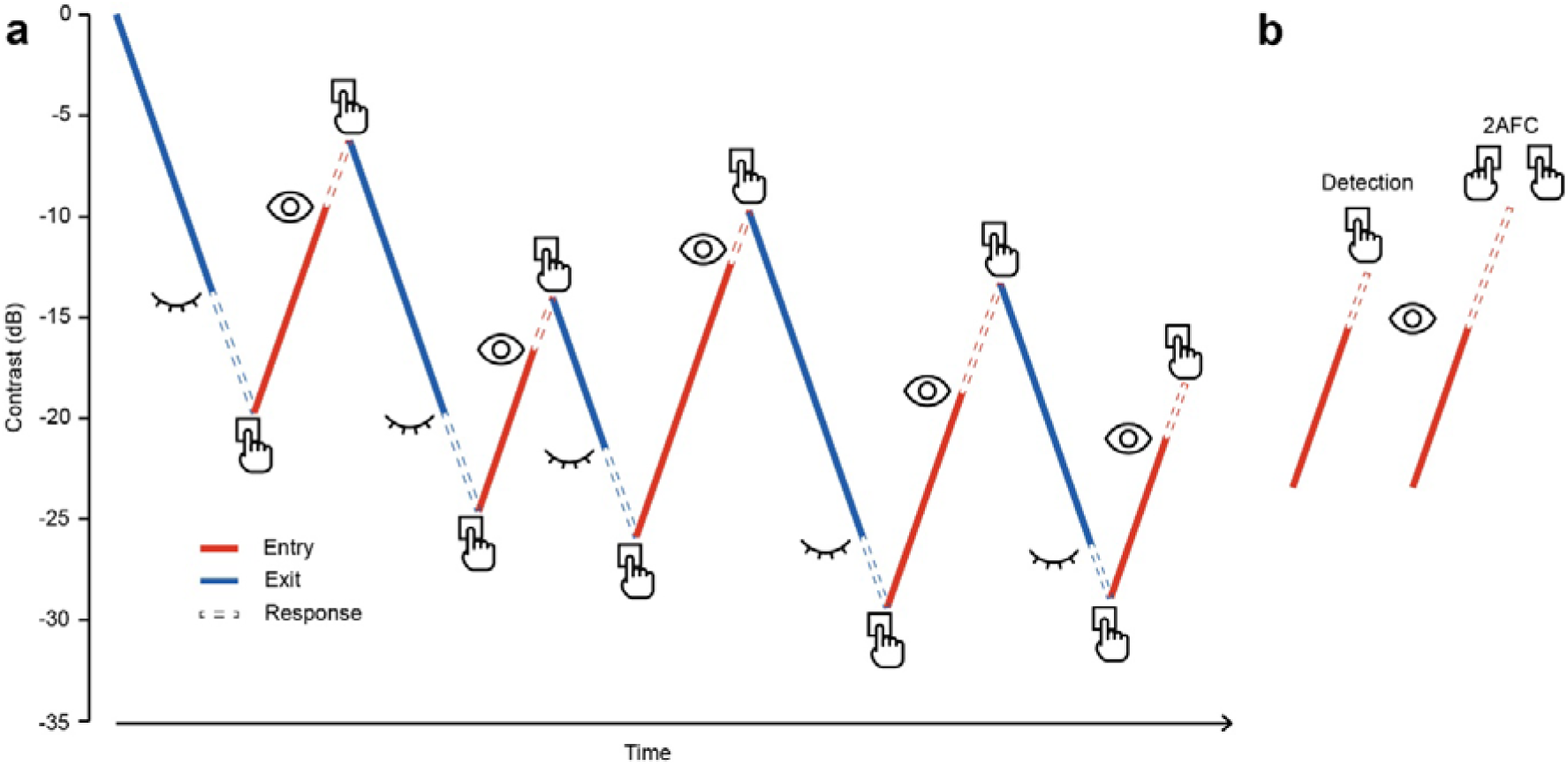
Hypothetical Components of tCFS Sequence. (**a**) Example of a tCFS sequence comprising five contrast ramping cycles. As indicated by the solid lines, target contrast continues to rise or fall until the observer reports that it enters (red) or exits (blue) consciousness. Hypothetical response-time components of each increasing and decreasing contrast ramp are presented as open dashed lines to illustrate “the overshoot problem”. (**b**) Example of increasing contrast ramps used in bCFS for the two different perceptual response tasks: a simple target detection, and a discrimination of target orientation (two-alternative, forced-choice). Response times for two-alternative forced-choice responses are predicted to be longer than those for simple detection.

Attentional research shows that stimulus onsets are often observed to be detected more efficiently than offsets, despite the sensory level changes being equivalent in magnitude (Chua, 2013; Cole et al., 2003, 2004; Cole & Kuhn, 2010; Donaldson & Yamamoto, 2016; Yantis & Jonides, 1984). This onset primacy (Cole et al., 2004) has been attributed to the evolutionary need for rapid detection of new stimuli to determine whether they pose a threat or not (Cole et al., 2003; Donaldson & Yamamoto, 2016). For an object already in awareness (as long as it does not present a danger), it is less consequential if it leaves awareness. Therefore, “the overshoot problem” may be smaller for the entry and exit of stimuli from awareness.

The purpose of this study is to address the overshoot problem by quantifying the reaction-time components for both breakthrough and suppression thresholds. We introduce a mathematical model that partials out the overshoot component to produce more accurate breakthrough and suppression thresholds and demonstrate that the model can be applied retrospectively to data from previously published CFS studies. The model works for simple target detection and for the slightly slower two-alternative target discrimination task. This latter point is important because existing tCFS studies (Alais et al., 2024, 2025; Alais & Kim, 2025; Coorey et al., 2026) have so far always used simple button presses to measure awareness thresholds rather than the discrimination task. This is because the tCFS paradigm emphasizes continuous tracking of perceptual state and a fast response is therefore preferable and because the slower discrimination task would exaggerate “the overshoot problem” (**Figure 1b**). The discrimination task, however, is widely used at breakthrough in many studies to validate that the target was seen and identified (Jiang et al., 2007; Stein, Hebart, et al., 2011; Sterzer et al., 2014; Zadbood et al., 2011) and so was incorporated into our tCFS experiments and modelling. Our data confirm a larger overshoot component for discrimination relative to the detection task and the model is able to quantify the reaction-time components for both. We also find the overshoot problem is asymmetrical, being greater for breakthrough compared to suppression thresholds. Both before and after reaction-time components are removed, targets require a greater decrease in contrast to exit awareness than the contrast increase required to enter awareness. Finally, our pupillometry data show that once the timing corrections are applied, pupil dilations that are known to occur when awareness of a target changes (Acquafredda et al., 2022; Privitera et al., 2010; Sperandio et al., 2018) are shown to align with the corrected time points.

## Methods

### Design

The study comprised two stimulus presentation paradigms, further divided by two response requirements, resulting in four tasks. The first of the presentation paradigms was the tCFS paradigm, during which contrast values when participants indicated the entrance and exit of target stimuli from visibility were measured. The second paradigm recorded response times to both the onset and offset of target stimuli. The responses for each task could be a simple detection response (i.e., a keypress response when target becomes visible or invisible) or a two-alternative forced choice orientation judgment (i.e., a specific keypress response for whether the target is oriented clockwise or counterclockwise upon becoming visible and a nonspecific keypress response when it becomes invisible).

For the tCFS task, participants completed four sequences, each comprising five contrast cycles (i.e., exit from visibility and re-entry into visibility) for each response requirement (**Figure 2a**). Every time participants indicated invisibility of the target stimulus, the target’s orientation could flip. As the target began at full contrast (i.e., RMS 100% or 0 dB) for each sequence, a sequence was composed of 6 stimuli. The repetitions (*n* = 12) of each orientation condition were balanced across the entirety of each response requirement. Overall, a 2 (*Response Requirement*: *Detection* vs *Orientation Judgment*) x 2 (*Visibility*: *Entry vs Exit) repeated-measures design was implemented with suppression depth (the difference between breakthrough and suppression thresholds)* as the dependent variable.

**Figure 2.**
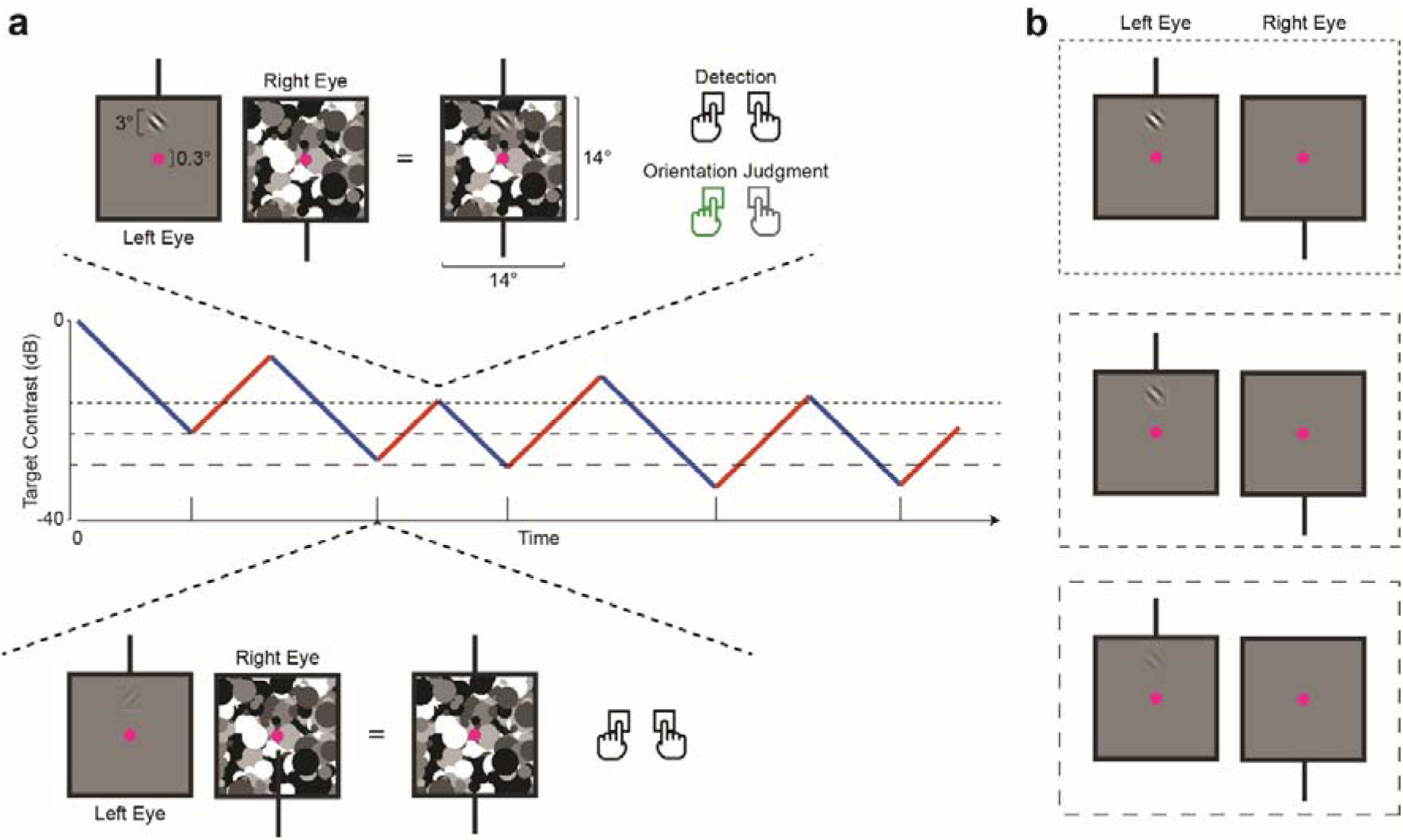
Schematic of Experimental Setup and Paradigm. (**a**) Example of a tCFS sequence comprising five contrast ramping cycles. Above is an example of the monocular input and the resultant binocular percept with responses illustrated for a breakthrough point. Below is an example of the monocular input and resultant binocular percept for a resuppression point. The horizontal dashed lines indicate the average breakthrough contrast, suppression contrast and the midpoint between the two; these contrast levels were used for the stimulus presentations in the response time task illustrated in (**b**). In these examples, the target was presented in the left eye while the mask was presented in the right; this was counterbalanced between participants.

For the response time task, participants’ response times to the onset and offset of target stimuli were measured. Participants completed twelve 15-target presentation sequences. The contrast of the presented targets took one of three values. The average contrast value at which target stimuli entered visibility, exited from visibility, and the midpoint between these values was calculated from the participants’ performance during the respective tCFS task block and used as contrast levels for the target presentations (**Figure 2b**). The orientation condition of the target stimuli was balanced across all repetitions and levels of presentation contrasts, resulting in 90 targets in each orientation condition. Overall, a 2 (*Response Requirement*: *Detection* vs *Orientation Judgment*) x 2 (*Visibility*: *Entry* vs *Exit*) x 3 (*Contrast*: *Entry*, *Midpoint*, *Exit*) repeated measures design was implemented with response time as the dependent variable. The *Entry* and *Exit* levels of the *Visibility* variable are synonymous with the onset and offset of the target stimuli.

The eye which received the Mondrian mask input was counterbalanced between participants. Due to the requirement to complete the tCFS tasks before the corresponding response time tasks, there were 6 valid block orders which were also counterbalanced. As a result, there were at least two participants for each combination of Mondrian input eye and block order, prior to exclusions.

### Participants

The sample consisted of a total of 26 third-year psychology students who participated in exchange for partial course credit. One participant was excluded from the current study due to having a condition that rendered correction to normal vision not possible. Two datasets were excluded due to incomplete participation. The final sample size was 23 (*M*_age_ ± *SD*_age_ = 21.70 years ± 2.75, 21 right-handed, 18 female). Each participant provided informed written consent before commencing and those not excluded from analysis had normal or corrected-to-normal vision. This study was approved by the University of Sydney Human Research Ethics Committee (HREC 2021/048). Each participant completed the experiment in a single one-hour session with approximately 30 min for task completion, excluding breaks.

### Apparatus

An HTC Vive Pro Eye head-mounted display (HMD) was used to display the virtual environment and a standard full format QWERTY keyboard was used for collecting responses. The HMD contained dual 3.5-inch high-resolution OLED displays (1440 x 1600 pixel resolution, 90 Hz refresh rate) with 110-degree field of view. The SRanipal Runtime (Version 1.3.2.0; SDK 1.3.6.8) was used to handle eye tracking data collection by the integrated Tobii eye-tracker within the HMD, sampled at 90 Hz.

The virtual reality (VR) environment and experimental procedure were designed and presented within Unity (Version 2020.3.14f1), using the SteamVR Plugin (Version 2.7.3; SDK 1.14.15) on a Windows 11 PC (2 × 16 GB, DDR5 4400 MHz) with a 12th Gen Intel Core i7-12700K processor (3.60 GHz), and an NVIDIA GeForce RTX 3070 (8 GB, GDDR6) graphics card. The VR environment was head-fixed, consisting of a mid-grey space (RGBA 128, 128, 128, 255; 0.5, 0.5, 0.5, 1.0).

### Stimuli

As per the tCFS (Alais et al., 2024) paradigm, each eye received different image inputs to facilitate interocular suppression. To one eye, an achromatic square Mondrian pattern with an RMS contrast fixed at 12%, spanning 14° of visual angle, was displayed. To the other eye, the displayed target stimuli were achromatic gratings tilted either 45° clockwise or counterclockwise from vertical with spatial frequencies of 1 cycle/° and diameters of 3° visual angle; the edges of the gratings were scaled with a raised cosine function to avoid distinct contours. A magenta fixation dot (diameter: 0.3° in visual angle) was displayed centrally for the entire duration of the experimental session. The target stimulus always appeared directly above the fixation point by 3.5° of visual angle. The rate of contrast change for the target stimuli was always 4.2 dB/s (i.e., 0.0467 dB/frame), ranging from 0 dB to - 39.993 dB, resulting in 858 discrete contrast values. Thus, at the 90 Hz frame rate of the HMD, the full contrast range was traversed in 9.53 s, much slower than the ∼1 s contrast ramps employed in bCFS. Note also that ramping contrast logarithmically (i.e, in dB) effectively linearises the response of early cortical neurons (Albrecht & Hamilton, 1982) so that the ramp is effective across its entire range (if contrast is ramped up linearly, most of the neural response to contrast change occurs in the early portion of the ramp and then saturates).

To ensure stable binocular fusion, a black marker was used during both calibration and testing as a fusion frame. It consisted of two aligned black boxes, each containing a vertical line segment, one pointing upward and the other downward. During the calibration phase, participants adjusted the position of the two monocular displays until they perceived a single black box with a partially occluded line running from top to bottom. The same marker remained on screen throughout the experiment to help maintain stable fusion. Between trial sequences, the marker turned from black to red as an indication of sequence completion and optional break times.

### Procedure

Participants were provided with a participant information statement, followed by an opportunity to ask any questions before they completed the consent form. All participants were informed that they could revoke their consent and participation at any time during the procedure if they wished not to provide their data or did not wish to complete their participation without the need to provide a reason. They were then introduced to the VR HMD and provided a verbal briefing of each task and their response requirements, including a thorough description of the input their individual eyes would receive and the overall percept they could expect. During this briefing, they were prompted for confirmation of understanding or for questions they may have had.

Once in VR, participants completed a 4-point calibration process for the in-built eye tracking component of the HMD, handled by the eye tracking software. Additional calibration was performed after any potential events that may have affected the accuracy of the tracking (e.g., signal drop-outs, breaks). The participants were then shown the experimental stimuli (see **Stimuli**), barring the Mondrian mask and target grating. They then calibrated the individual eye displays using the up and down arrow keys, either moving the displays in converging (i.e., left eye input shifting right while right eye input shifted left) or diverging directions (i.e., left eye input shifting left while right eye input shifted right), until they perceived a binocular percept of a square occluding a line running vertically behind and a singular magenta fixation dot at the centre. Participants were then allowed to experience the components of the tCFS paradigm (i.e. dynamic Mondrian mask and contrast ramping of target grating stimuli), without requirement for proper responses, both monocularly and in binocular. Upon confirmation of their understanding and satisfaction, they were provided with instructions for their first task and began the respective practice sequences. Before the beginning of every one of the four task blocks, the participants underwent a practice sequence and had the option of repeating this practice sequence until they were comfortable completing the task. Most individuals only completed a single practice sequence for each of the tasks with the maximum repetitions being three. Participants were informed that breaks could be taken during periods when the fusion markers were red and that, to initiate the subsequent sequence, they should press either response key (i.e., “Q” or “P”).

#### Tracking Continuous Flash Suppression

For the tCFS tasks, the participants were instructed to report the moment when the visibility of the target stimuli changed between visible and invisible. They were explicitly told to report when targets became visible enough to perceive the “tilted lines” and when they only perceived the mask. For the block when simple detection responses were required, participants could respond by pressing either the “Q” or “P” keys to indicate both visibility and invisibility. During the orientation judgment block, the participants were instructed to press “Q” if the target was counterclockwise rotated or “P” if clockwise rotated when it entered visibility and either key when it exited. When participants indicated confusion or it was made apparent the task was misunderstood from the practice sequences, alternative verbal instructions were provided. Such instructions included “Press Q when the target is tilted to the left” or “Press P when the lines are pointing to North East, not North West”, amongst many others.

#### On-Off Response Time

For the response time tasks, the participants were instructed to respond as quickly as possible when any target stimuli appeared or disappeared. For the block when simple detection responses were required, participants could respond by pressing either the “Q” or “P” keys to indicate whenever the targets appeared or disappeared. During the orientation judgment block, the participants were instructed to press “Q” if the target was counterclockwise rotated or “P” if clockwise rotated when it appeared and either key when it disappeared. When participants indicated confusion or it was made apparent the task was misunderstood from the practice sequences, alternative verbal instructions were provided as outlined previously.

### Exclusions

Beyond the participant exclusions mentioned above, there were exclusions of individual data points applied before analysis.

Given the positive skew typical of response time data, which was also present in the current study, response times exceeding 3 standard deviations within each participant’s data were removed as outliers. This exclusion approach has been shown to minimise biases that may occur as a result of outlier removal while limiting the data removed (Berger & Kiefer, 2021).

As for the data from the tCFS tasks, any suppression depth values that end at or beyond the extremes of the contrast range were excluded for ceiling (≥ 0 dB) or floor (≤ -39.993 dB) effects. These exclusion criteria were applied after the corrective model was applied.

### Statistical Analysis

#### Response Times

Due to the positive skew found in response time data, a log transformation was applied to the data before being fitted with a linear mixed model. A linear mixed model was chosen over a repeated-measures ANOVA as it was better suited to estimate the effect of the target contrast, a continuous variable. The equation of the linear model was as follows:

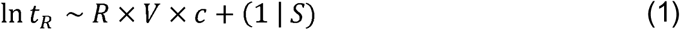

where the dependent variable, the natural log of response times *t_R_*, is the linear combination of the fixed effects of the binary *Response Requirement* variable *R*, binary *Visibility* variable of the target *V*, and the continuous variable of target contrast *c*, including the possibility of interaction between such variables and accommodating for a random intercept for each participant *S*.

#### Suppression Depth

According to the method of calculation employed by Alais et al. (2024), suppression depth is the difference between the average contrast values, in decibels, at which participants indicated the target stimulus entered visibility and when it exited visibility (note that being on a log scale, this contrast difference is in effect a ratio of contrasts). However, to address the aims of the current study, depth was instead calculated as the difference between consecutive contrast values where targets exited and entered visibility, thereby alternating between positive and negative depths. The absolute value of each depth was taken to facilitate comparison between depths. The initial decrease in contrast of every tCFS sequence was excluded from analyses as the initiation at full contrast would overinflate the magnitude of that depth.

A three-way 2 (*Response Requirement*: *Detection* vs *Orientation Judgment*) x 2 (*Visibility*: *Entry* vs *Exit*) x 2 (*Correction*: *Before* vs *After*) repeated-measures ANOVA was performed on the magnitude (i.e., absolute value) of suppression depth, before and after the correction model was applied.

#### Pupillometry

As changes in pupil diameter have been correlated with task demands, optical stimulation, and arousal, pupil diameter is a promising physiological marker of interocular suppression. To investigate whether transitions to and from conscious awareness could be observed physiologically, pupil diameter during awareness and suppression of the target stimuli was analysed. Specifically, whether pupil diameter shows changes around corrected points of breakthrough and suppression was of interest.

Pupil data for both target- and mask-presented eyes were segmented by breakthrough and suppression points. Any segment containing a blink was removed to avoid any transient changes induced by changes in optical stimulation. Remaining segments were demeaned and processed with a Savitzky–Golay filter (20 samples, 0.222 s, 2^nd^ order). Due to the variable length of time between breakthrough and suppression points and the difference in the correction across tasks and transitions, these segments were only comparable after normalization to percentiles (**Figure S1**). For the same reasons, the data were epoched around each corrected breakthrough and suppression point. These epochs extended from –0.5 to 0.5 s from the corrected points. To determine the significance of the corrected breakthrough and suppression points in changes in pupil diameter, a linear mixed model was fitted:

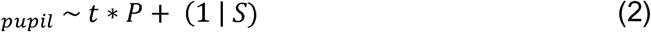

where the dependent variable, the demeaned pupil diameter ⌀*_pupil_*, is the linear combination of the fixed effects of the binary *Phase* variable *P*, and the continuous variable of *Time t*, with a random intercept for each participant *S*. *Phase* was whether the time point was before or after the corrected breakthrough or suppression point. This model approach is analogous to a breakpoint, or segmented, regression with an informed breakpoint that accounts for individual differences in mean pupil diameter. An equivalent analysis was performed on epochs centred on reported suppression and breakthrough points (**Figure S2**).

### Suppression Depth Correction Model

From the linear model, described in (1), the main effects of *Response Requirement*, *Visibility* and *Contrast*, and all of their two- and three-way interactions on log-transformed response times were statistically significant. As such, to remove the influence of the response-making process from suppression depth and therefore, provide a more accurate reflection of perceptual awareness, any correction model must consider all these factors.

First, the linear model was broken down into a more applicable form. Since the only continuous variable in (1) was *Contrast*, it can be viewed as a simple linear regression with the contrast value of the targets as the predictor, as shown here:

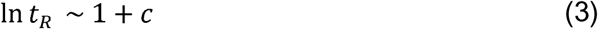

but applied separately for each of the conditions that result from the combination of *Response Requirement* and *Visibility* for each participant. From this, for ease of implementation, we reverse the log transformation of the response times, giving us:

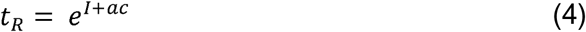

where *a* is the fitted slope estimate, *I* is the fitted intercept estimate for the assumed linear relation between response time and target contrast for the specific condition of each participant.

Because contrast changed at a constant rate over time, the difference between when the target entered and exited visibility can be interchangeably expressed as the difference between the contrast values (as per the suppression depth calculation) or as a difference in time Δ*t*:

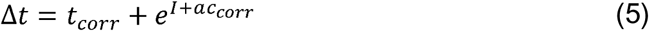

where *t_corr_* is the hypothetical (corrected) time point at which the target either entered or exited visibility and *c_corr_* is the corresponding contrast at this time point.

We can convert the hypothetical contrast value at which the target either entered visibility or exited it, using the following equation:

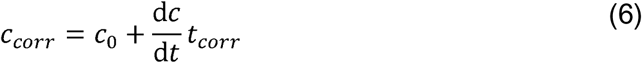

where *c*_0_ is the initial contrast that the target was changing from and 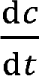 is the directional rate at which the contrast was changing. This equation can then be substituted into (5) to produce:

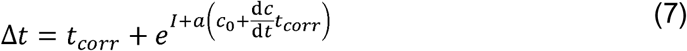

solving this for *t_corr_*:

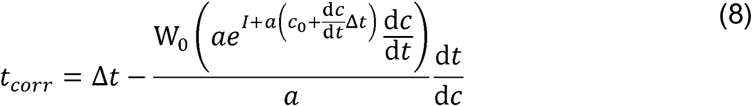

where W_0_is the principal branch of the Lambert W function. Multiply the corrected change in time by the rate of contrast change to get the corrected suppression depth *d_corr_*:

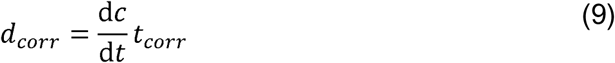

## Results

### Response Time

A Linear Mixed Model was fitted for the natural logarithm of response times with the fixed effect variables consisting of the binary *Response Requirement* variable, *R*, the binary *Visibility* variable, *V*, and the continuous *Contrast* variable, *c*. The *Contrast* variable was mean-centred before the model was fitted. The model estimated random intercepts for each participant. The equation for the fixed effects model was the following:

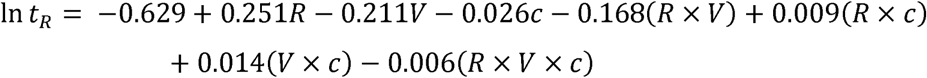

where *Response Requirement* had the *Detection* task and *Visibility* had *Entry* as the reference/baseline levels.

When participants were just detecting the onset of a target, the natural logarithm of response times decreased on average by .026 units (*b* = -.026, *SE_b_* = .001, *z* = -25.442, *p* < .001, 95% CI [-.028, -.024]) for every decibel increase in target contrast (**Figure 3a**). As for detecting the offset of targets, this relationship became weaker, such that for every decibel increase in contrast the natural logarithm of response times decreased on average by .014 units less (*b* = .014, *SE_b_* = .001, *z* = 10.004, *p* < .001, 95% CI [.011, .017]).

**Figure 3.**
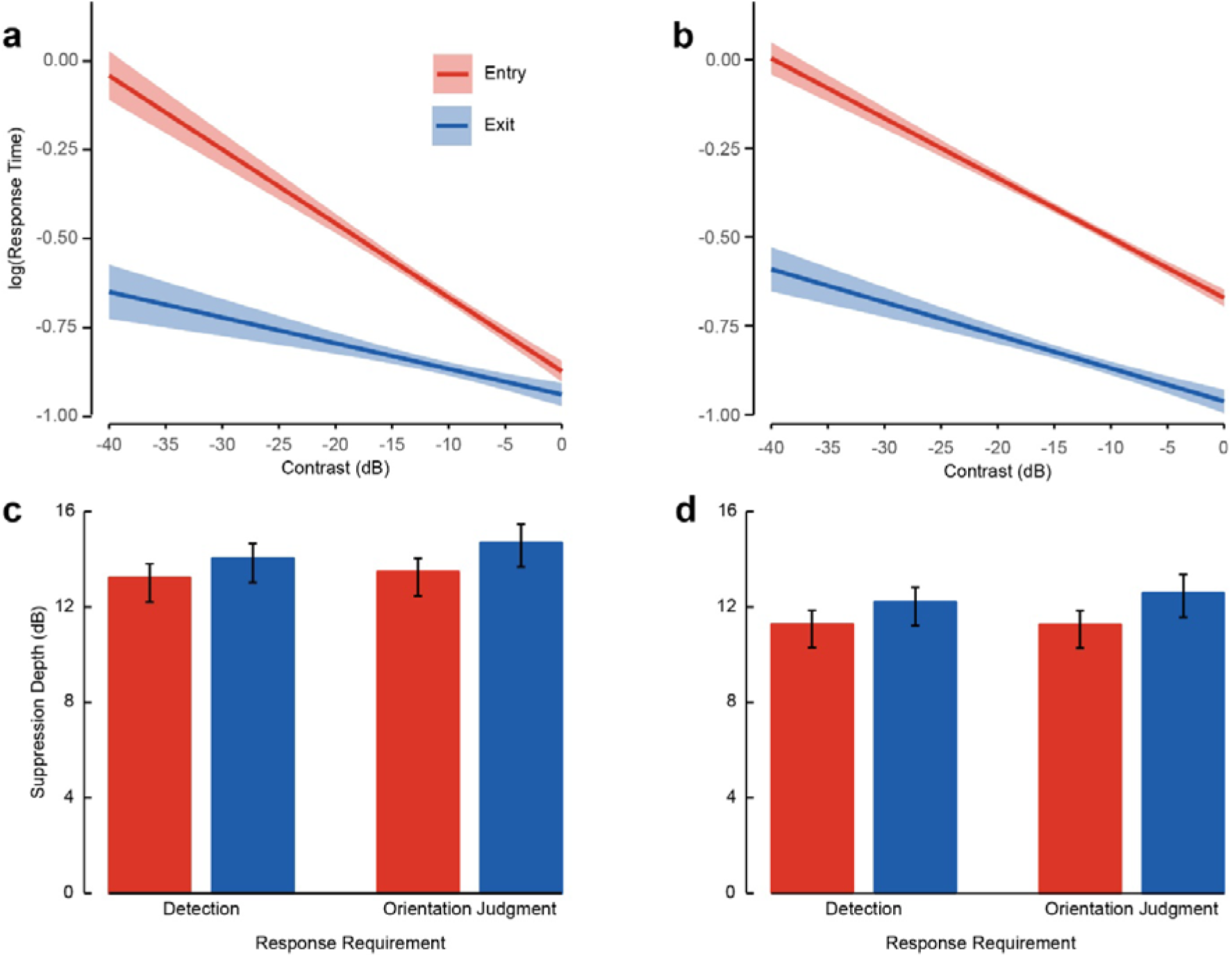
Response Time and Suppression Depth from Simple Detection and Orientation Judgment. (**a**) Displays the estimated fixed effects of responding at *Entry* and *Exit* of *Visibility* over target contrast for natural logarithm of simple detection response times. (**b**) Displays the estimated fixed effects of responding at *Entry* and *Exit* of *Visibility* over target contrast for natural logarithm of two-alternative forced choice orientation judgment response times. Bar plots display the effect of different response requirements at *Entry* and *Exit* of *Visibility* on suppression depth (**c**) before and (**d**) after applying the correction for response time. Shaded regions and error bars represent 95% confidence intervals.

The natural logarithm of response times increased on average by .251 units (*b* = .251, *SE_b_* = .010, *z* = 24.011, *p* < .001, 95% CI [.230, .271]) when the responses were orientation judgments as opposed to simple detection to target onsets, averaged over contrast (**Figure 3b**). This positive relationship was strengthened on average by .009 units (*b* = .009, *SE_b_* = .001, *z* = 6.516, *p* < .001, 95% CI [.006, .012]) for each decibel increase in target contrast.

Averaged over contrast, when participants were detecting the offset of targets as opposed to the onsets, the natural logarithm of response times was on average lower by .211 units (*b* = -.211, *SE_b_* = .010, *z* = -20.372, *p* < .001, 95% CI [-.231, -.191]). This difference between onset and offset responses increased by .168 units (*b* = -.168, *SE_b_* = .015, *z* = -11.436, *p* < .001, 95% CI [-.197, -.139]) when the participants were making orientation judgments at onset instead. Additionally, this increased difference became even larger by .006 units (*b* = -.006, *SE_b_* = .002, *z* = -3.343, *p* = .001, 95% CI [-.010, -.003]) for every decibel increase in target contrast.

### Suppression Depth

A three-way 2 (Response Requirement: Detection vs Orientation Judgment) × 2 (Visibility: Entry vs Exit) × 2 (Correction: Before vs After) repeated-measures ANOVA was run on suppression depth (**Figure 3c, d**). There was a main effect of *Correction*, such that suppression depths were significantly smaller after applying the correction model (η^2^ = .269, *F*(1, 22) = 268.6, *p* < .001). The suppression depth produced when targets were increasing in contrast and entering visibility was significantly smaller than when they were decreasing in contrast and exiting visibility (η^2^ = .063, *F*(1, 22) = 11.89, *p* = .002). There was a significant interaction between *Correction* and *Response Requirement*, such that the decrease in suppression depths before and after applying the correction model was larger for those calculated from the orientation judgment task than those from the simple detection task (η^2^ = .001, *F*(1, 22) = 7.48, *p* = .012). There was a significant interaction between *Correction* and *Visibility*, such that the contrast change during suppression was decreased more after correction than the contrast change when the target was not suppressed (η^2^ = .002, *F*(1, 22) = 7.78, *p* = .011). There was no significant main effect of *Response Requirement* and no other significant interaction effects, *p*’s > .05.

### Pupillometry

Linear Mixed Models were fitted for pupil diameter with the fixed effect variables consisting of the continuous *Time* variable, *t*, and the binary *Phase* variable, *P*. The *Time* variable was epoched to the corrected time of suppression or breakthrough from 500 ms before to 500 ms after. *Phase* was whether the time point was before or after the corrected breakthrough or suppression point. The model estimated random intercepts for each participant. Models were fitted for each eye for each *Visibility* condition. The equation for each fixed effects model was the following:

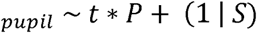

As the target was exiting conscious awareness, the pupils of both the target-presented eye (**Figure 4a**) and Mondrian mask-presented eye (**Figure 4b**) were constricting at the rate of .030 and .027 mm/s, respectively (target: *b* = -.030, *SE_b_* = .007, *z* = -3.984, *p* < .001, 95% CI [-.044, -.015]; Mondrian: *b* = -.027, *SE_b_* = .007, *z* = -3.867, *p* < .001, 95% CI [-.041, -.014]). After the corrected suppression time, the rate of change in the pupil diameter of the target eye was increased by .052 mm/s (*b* = .052, *SE_b_* = .011, *z* = 4.608, *p* < .001, 95% CI [.030, .075]), meaning the pupil was dilating instead of constricting (*b* = .019, *SE_b_* = .007, *z* = 2.800, *p* = .005, 95% CI [.006, .032]). Meanwhile, the pupil of the masked eye exhibited a change in rate of .027 mm/s (*b* = .027, *SE_b_* = .011, *z* = 2.521, *p* = .012, 95% CI [.006, .049]), no longer changing in diameter with time (*b* = .000, *SE_b_* = .006, *z* = 0.034, *p* = .973, 95% CI [-.012, .013]).

**Figure 4.**
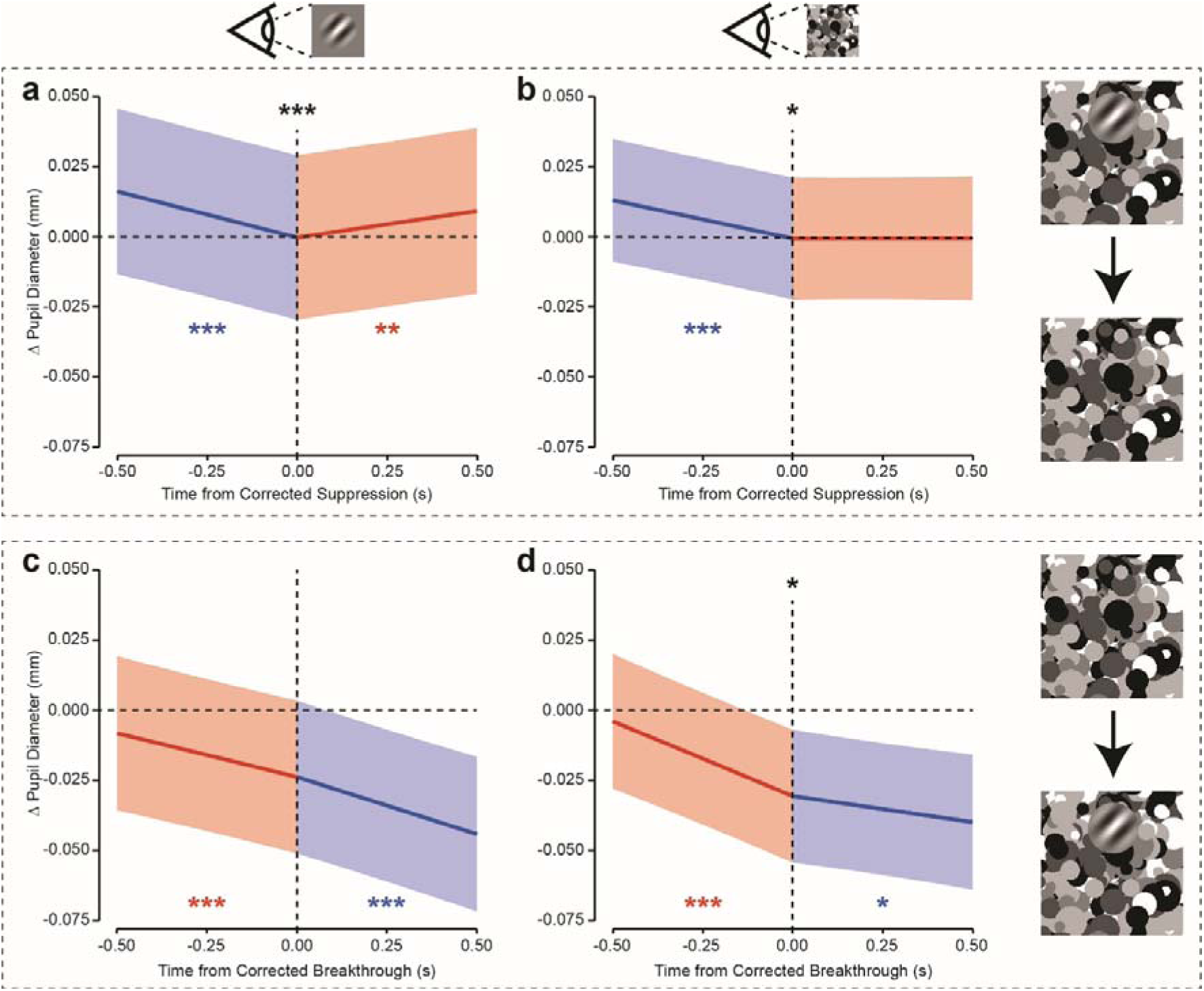
Pupil Diameter as a Function of Time Around Corrected Suppression and Breakthrough. (**a**) Displays the estimated fixed effect of time on pupil diameter of the target presented eye 500 ms before and after the corrected transition from conscious awareness to suppression of targets. (**b**) Same as (**a**), but for the eye presented with the Mondrian mask. (**c-d**) Same as (**a-b**), but for the corrected transition from suppression to conscious awareness. Coloured lines and regions correspond to the state of conscious awareness of the target; blue for targets within awareness and red for suppressed targets. Shaded regions represent 95% confidence intervals. \**p* < .05. \*\**p* < .01. \*\*\**p* < .001, where coloured significance indicators are for corresponding slopes against 0 and black indicators are for differences in slopes across transition point.

As the corrected point of breakthrough was being approached, the pupils of both the target-presented eye (**Figure 4c**) and Mondrian mask-presented eye (**Figure 4d**) were constricting at the rate of .027 and .054 mm/s, respectively (target: *b* = -.027, *SE_b_* = .010, *z* = -2.747, *p* = .006, 95% CI [-.047, -.008]; Mondrian: *b* = -.054, *SE_b_* = .010, *z* = -5.180, *p* < .001, 95% CI [-.074, -.033]). After the corrected breakthrough, the rate of the change in the target eye did not differ (*b* = -.009, *SE_b_* = .014, *z* = -0.639, *p* = .523, 95% CI [-.038, .019]) while the constriction of the masked eye pupil slowed by .035 mm/s (*b* = .035, *SE_b_* = .015, *z* = 2.310, *p* = .021, 95% CI [.005, .064]) to .019 mm/s (*b* = -.019, *SE_b_* = .009, *z* = -2.145, *p* = .032, 95% CI [-.035, -.002]).

### Application to Previous Datasets

For ease of interpretation, the model equation can be resolved for any of the conditions resulting from combination of the binary variables, *Response Requirement* and *Visibility* (**Figure 3a, b**). The resolved equations are as follows:

In *t_R_* = −0.629 − 0.026*c* when *Response Requirement* is *Detection* and *Visibility* is *Entry*

In *t_R_* = −0.840 − 0.012*c* when *Response Requirement* is *Detection* and *Visibility* is *Exit*

In *t_R_* = −0.378 − 0.017*c* when *Response Requirement* is *Orientation Judgment* and *Visibility* is *Entry*

In *t_R_* = −0.757 − 0.009*c* when *Response Requirement* is *Orientation Judgment* and *Visibility* is *Exit*

For bCFS studies and the ascending half of tCFS cycles, the first equation can be used in place of (3) for the application of the corrective model, resulting in the following:

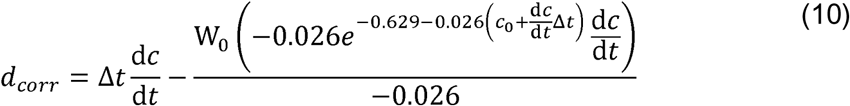

Δ*t* is the suppression depth time or breakthrough time, 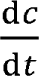 is the contrast ramp in dB/s specific to the study and *d_corr_* is the corrected suppression depth or breakthrough contrast in dB, either of which can be converted to time. It must be noted, however, that due to potential influences of individual differences, such as age, sex, and training, this simplified correction should only be applied to samples with similar demographics (i.e., undergraduate students).

For the descending half of tCFS cycles, the correction would be:

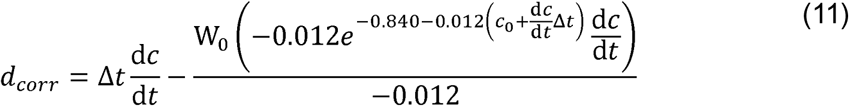

### bCFS – Linear Contrast Ramps

To provide a guide on applying the correction to classic bCFS studies, we will walk through the process on the data from the seminal bCFS study by Jiang et al. (2007). Using pixel extraction, the suppression time (i.e., breakthrough time) of each participant for upright and inverted faces was approximated (**Figure 5a**). Due to their use of a broken y-axis, the tenth participant’s data could not be estimated reliably and was therefore excluded from our re-analysis (**Figure 5b**).

**Figure 5.**
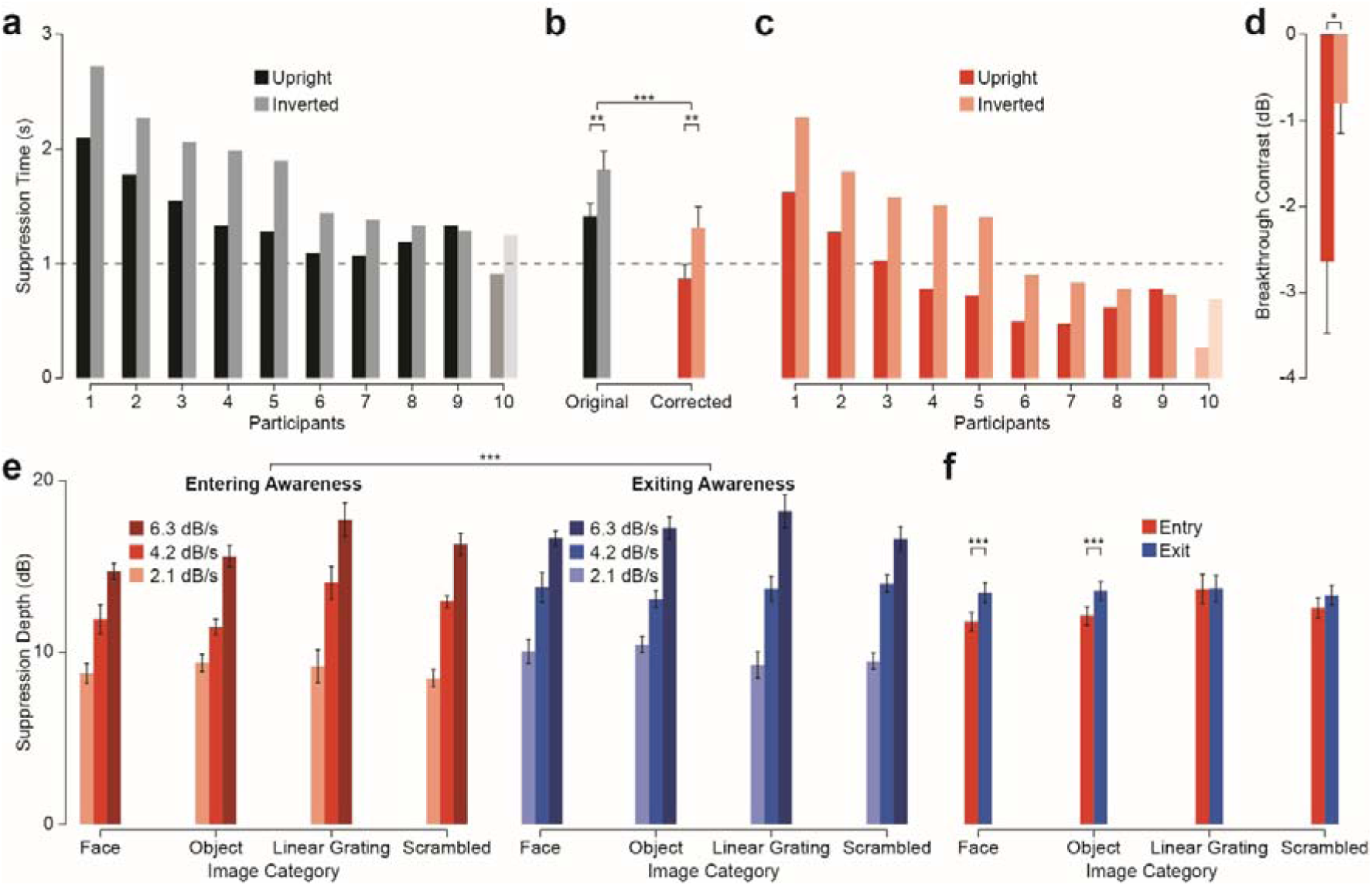
Pupil Diameter as a Function of Time Around Corrected Suppression and Breakthrough. (**a**) Suppression times, equivalent to breakthrough times, of each participant in Experiment 1 of Jiang et al.’s (2007) seminal bCFS study. Values were estimated using pixel extraction of their Figure 2. (**b**) Average suppression time across nine out of ten participants, excluding the tenth due to the ambiguity in broken axis in original figure. (**c**) Same as (**a**), but after correcting for response times after applying correction model. (**d**) Same as (**b**), but only for corrected values after conversion to decibels. (**e**) Suppression depth values from Experiment 3 of Alais et al.’s (2024) seminal tCFS study after application of the response time correction model. (**f**) Values in (**e**), but averaged across target contrast ramp speeds.

In their study, Jiang et al. (2007) ramped their targets in linear contrast from 0 to 1 in 1 s, resulting in a non-constant decibel contrast rate (see **Stimuli**). First, the linear contrast must be converted to decibels:

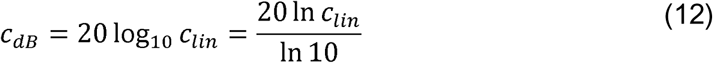

Substitute this into the exponent seen within the Lambert W function (10), it resolves to be:

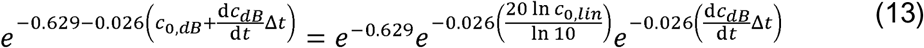

where the exponent concerning starting contrast simplifies to:

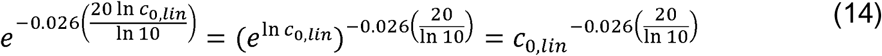

Let us simplify the following equations by establishing:

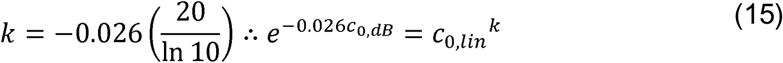

Moving onto the contrast ramp, conversion renders the decibel rate a variable linear rate:

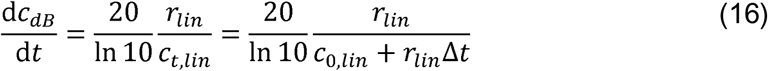

The contrast change during breakthrough times will then be:

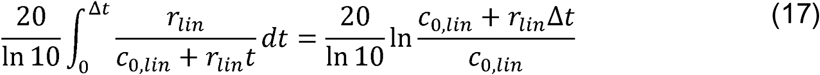

which would convert the exponent concerning contrast change to:

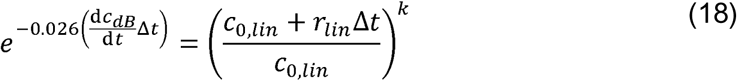

therefore, the exponent in the Lambert W function becomes:

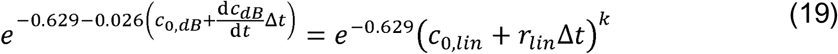

Finally, the correction model becomes the following for linear contrast ramping:

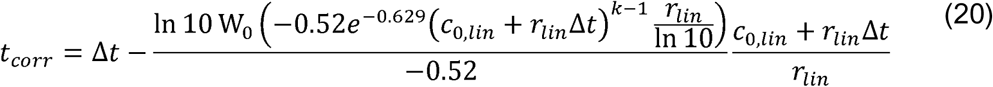

Jiang et al. (2007) used a linear contrast rate *r_lin_* of 1 Hz and had beginning contrasts *c*_0,*lin*_ of 0, which substitutes into (20) to become:

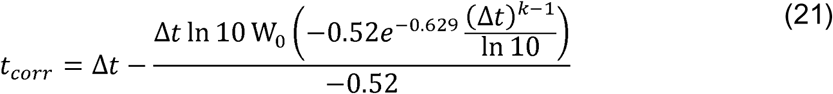

A two-way 2 (Correction: Before vs After) × 2 (Face: Upright vs Inverted) repeated-measures ANOVA was run on suppression times (**Figure 5a-c**). As expected of a corrective removal of response time estimates, suppression times were significantly smaller after applying the linear ramp correction model (η^2^ = .535, *F*(1, 8) = 1594, *p* < .001; **Figure 5c**). Confirming Jiang et al.’s (2007) finding of an upright face priority advantage, suppression times were longer for inverted face images (η^2^ = .355, *F*(1, 8) = 26.78, *p* < .001). There was a significant interaction between *Correction* and *Face*, such that the advantage for upright faces was larger after correcting for response times (η^2^ < .001, *F*(1, 8) = 27.98, *p* < .001). This reflects the response time scaling with target contrast; higher contrast targets take shorter times to respond to and hence the overshoot component removed for inverted face targets was smaller than for upright face targets. Converting the suppression times to breakthrough contrasts, the upright face advantage was retained with breakthrough occurring at lower contrasts than for inverted faces (Cohen’s *d* = 0.849, *t*(8) = 2.548, *p* = .034; **Figure 5d**).

### tCFS – Logarithmic Contrast Ramps

We additionally re-analysed data from the third experiment of the seminal tCFS study by Alais et al. (2024). This experiment was chosen as the target contrast ramp was varied as a factor and multiple image categories were used as target images. The correction model (10 and 11) needed no alteration as contrast was ramped on a decibel scale. Due to one participant consistently letting targets reach minimum contrast (i.e., -33.98 dB), we excluded an additional participant on top of the original singular exclusion by Alais et al. (2024), resulting in 15 participants’ data remaining for analysis.

A three-way 2 (Visibility: Entry vs Exit) × 2 (Image Category: 2, 7, Linear Grating, and Scrambled) × 3 (Ramp: 2.1, 4.2, and 6.3 dB/s) repeated-measures ANOVA was run on corrected suppression depth (**Figure 5d**). The suppression depth produced when targets were increasing in contrast and entering visibility was significantly smaller than when they were decreasing in contrast and exiting visibility (η^2^ = .015, *F*(1, 14) = 17.31, *p* < .001). There was no significant main effect of *Image Category* (η^2^ = .010, *F*(1, 14) = 1.073, *p* = .371). Mauchly’s test of sphericity indicated a violation of sphericity for the main effect of *Ramp* and the *Visibility × Image Category* interaction. We applied Greenhouse-Geisser corrections for these effects. There was a significant main effect of *Ramp*, but no significant interaction involving *Ramp* (*p*’s > .05), such that suppression depth measured during both target awareness and unawareness similarly varied with target contrast ramp speed (η^2^ = .549, *F*(1.451, 20.31) = 117.1, *p* < .001). The *Visibility × Image Category* interaction was significant, such that the difference between suppression depth during target awareness and unawareness differed across image categories (η^2^ = .006, *F*(1.562, 21.87) = 8.416, *p* = .003; **Figure 5f**). This interaction appears to be driven by the fact that there is no significant difference between suppression depth measures for linear grating images (Cohen’s *d* = 0.008, *t*(14) = 0.082, *p_holm_* = 1.000) and for scrambled images (Cohen’s *d* = 0.148, *t*(14) = 2.837, *p_holm_* = .316) while the other image categories had higher suppression depth during visibility (2: Cohen’s *d* = 0.335, *t*(14) = 7.222, *p_holm_* < .001; 7: Cohen’s *d* = 0.287, *t*(14) = 5.725, *p_holm_* = .001). There were no other significant interaction effects, *p*’s > .05.

## Discussion

This study aimed to quantify “the overshoot problem” in estimating CFS thresholds that arises from the observer’s reaction time to respond to the target while it continues to ramp up (or down) in contrast. Measuring these reaction-time components revealed that overshoot is asymmetrical, being greater for stimuli entering awareness (targets rising in contrast towards breakthrough threshold) than for exiting awareness (visible targets falling in contrast towards suppression threshold). We introduced a mathematical model that quantifies the overshoot component so that it can be partialled out of CFS threshold estimates to provide more accurate breakthrough and suppression thresholds.

The problem of overshoot is inherent in any CFS method that ramps the target’s contrast over time, such as bCFS and tCFS. This is particularly true when the contrast ramp is steep, as is often the case in bCFS studies where the stimulus increases from minimum to maximum in just a second or so (many times steeper than used in the tCFS method). For any given response time, a rapid contrast change inevitably yields a threshold with a greater overshoot component than a slower contrast change. Consider an observer with a response time of ∼300 ms who does a bCFS task in which the target ramps from minimum (0) to maximum (1.0) in 1 second. Supposing their true contrast breakthrough threshold were 0.60, their measured threshold (given a 300 ms RT) would be 0.90, a staggering 50% higher than their true threshold. The threshold for the same observer and RT repeating the experiment with a 4-second contrast ramp would yield a threshold of 0.675. Not only does this problem conflate threshold and ramp-rate in a given study, but comparisons with other studies would reveal spurious threshold differences for the same stimuli if a different ramp rate were used.

Overshoot is less of a problem for tCFS, where ramps are more gradual (e.g., total ramp time from min to max in Alais et al., 2024 was ∼8 s) and for other bCFS studies that used gradual ramps. However, because ramp timing is not standardised it is a variable that adds noise to breakthrough and suppression thresholds and hinders meaningful comparisons between studies. The corrective model we introduce here is very useful in this respect because it can be applied to previous and future studies to take into account the effect of contrast ramp time and response time and it can be applied to breakthrough and suppression thresholds. The model is based on logarithmic contrast change over time but can be adapted to deal with data from linear contrast ramps and will partial out the threshold component due to overshoot to reveal more accurate threshold estimates for breakthrough or suppression (see **Figure 1**). This could prove very useful in the CFS field where a number of controversies have emerged around the issues of visual processing outside of awareness and privileged access to awareness for certain kinds of images. A systematic comparison after overshoot correction of the key studies in these debates may help resolve them and advance the field.

Another factor that affects response time and thus contributes to “the overshoot problem” is target contrast at breakthrough. The top panels of **Figure 3** show that response times are not fixed as a function of target contrast. Instead, there is a significant increase of response times as the target contrast at breakthrough reduces. Breakthrough thresholds therefore can be expected to vary significantly among published studies based on how strong their Mondrian mask was. A high-contrast mask would easily suppress a low-contrast target, meaning the target would need to reach a higher contrast to achieve breakthrough – which would be reported with a relatively fast response time (**Figure 3a, b**). For a low-contrast mask, however, target breakthrough would be reached at a lower contrast and consequently with a relatively slower response time. For this reason, careful consideration should be given to mask contrast in CFS studies and, again, this may be a factor that contributes to controversy in the field by adding unnecessary variability among different labs.

One of our predictions was that stimulus onsets (targets breaking into awareness) would elicit shorter response times than stimulus offsets (visible targets succumbing to suppression). This was based on ‘onset primacy’, the finding that onset of a novel stimulus is more relevant than the offset of a known stimulus. Contrary to the expected onset primacy effect, we found that the response time for targets entering awareness was longer than for targets exiting awareness. Interestingly, this asymmetry was observed regardless of whether the task was simply to detect the target’s breakthrough or discriminate its orientation at breakthrough. A possible explanation for the increased response times for onsets may be that the set of presented targets quickly became familiar and eroded stimulus novelty (Cole et al., 2004). It is also possible that continually ramping the target’s contrast introduced weaker or ambiguous mental templates that increased the difficulty of template matching and led to increased response times (Lleras et al., 2020). There may also be a neurophysiological basis for longer response times at breakthrough as it has been shown that neural response latencies in primate visual cortex are shorter for stimulus offsets than stimulus onsets (Bair, 2004; Bair et al., 2002).

Another important methodological implication of our findings is that two-alternative, forced-choice tasks can be implemented in tCFS designs without impacting the validity of its threshold estimates. The appeal of tCFS is that the trial is continuous and data collection is rapid: the target simply ramps up and down in contrast, bouncing continuously between two thresholds (Alais et al., 2024), and the participant simply tracks their perceptual state and presses a key every time it changes (producing the reversals seen in **Figure 1**). The tCFS studies published to date used a simple task of detecting the appearance or disappearance of the target to avoid long response times interrupting the flow of the experiment. Now that response times can be quantified and partialled out of threshold estimates, discrimination tasks can be used in tCFS. Discrimination tasks allow probing of the observer’s conscious perception of a stimulus (detection may simply indicate the onset of ‘something’ but not a clear percept which can be interrogated) and thus verification of their experience. This provides an avenue to explore more ecologically applicable experiences beyond simple awareness. Furthermore, researchers can simply take the parameters of the (log-linear) relationship between target contrast and response time for any response task and substitute them into the corrective model we have proposed to correct for “the overshoot problem” for their particular task and circumstances.

In the current data set, after correcting for “the overshoot problem”, suppression depth did not differ between the simple detection task and the orientation discrimination task. Let us assume that the time between breakthrough and suppression is made up of two components: the time the target is under interocular suppression (if it is ramping up) or resisting suppression (if it is ramping down) and the observer’s response time. In removing the contrast change that occurred during the response component, we have effectively isolated the contrast change required for the transition between conscious awareness and unawareness. The lack of threshold difference after correction for the two types of tasks indicates a similar degree of suppression, and resistance to suppression, for each task. This is as it should be because the stimuli for both response tasks were the same and would be expected to produce identical interocular dominance and suppression (Brascamp et al., 2015). In observing the same suppression depths between the tasks after removal of the response time overshoot, the transition from and into interocular suppression appears to occur without the influence of higher-level context (i.e., the task of discriminating the target stimulus). A future study could employ neurophysiological measures (e.g., EEG, fMRI) to identify the point when a target enters awareness, indicated by changes in evoked potentials beyond those caused by contrast change (Kurita et al., 2021; Metzger et al., 2017; Pitts & Britz, 2011; Shinozaki & Takeda, 2008), and when the target’s features are identifiable (i.e., when decodability of its neural representation exceeds chance (Contini et al., 2017; Misaki et al., 2010; Tang et al., 2019).

One interesting observation was that the difference between breakthrough and suppression thresholds depended on the direction of contrast change. When the target was suppressed but ramping up towards awareness, suppression depth was less than when the target was visible but ramping down towards suppression. This increased “suppression resistance” persisted after correcting for response time. When a target is suppressed, there are limited factors that could promote it into awareness, particularly as CFS produces very deep suppression (several times stronger than binocular rivalry suppression: Alais et al., 2024; Blake & Camisa, 1979). According to Levelt’s propositions (Brascamp et al., 2015), as a stimulus increases in contrast, its relative predominance against a constant interocular rival increases. In bCFS and tCFS, the target ramps up in contrast and at breakthrough, neurons representing the target eventually dominate those representing the mask (Economides et al., 2012; Paffen et al., 2008). This is a simple, low-level account with limited scope for higher-level effects: if target activity is upregulated or baseline activity is decreased, the target’s ability to overcome suppression from the mask increases (Friston, 2005; Lee & Mumford, 2003; Spratling, 2008). However, once the target breaks into awareness, additional factors may come into play and may endow it with suppression resistance. One potential contributor to sustaining awareness of a target as it declines towards suppression would be the attention directed toward it, which could be driven by its salience, task relevance, and higher-level processing requirements (Connor et al., 2004; Corbetta & Shulman, 2002; Paneri & Gregoriou, 2017). This was demonstrated in our re-analysis of Alais et al.’s (2024) results, where suppression resistance was greater than suppression depth for target images involving higher-level semantic processing (i.e., faces and familiar objects), but not for simplistic stimuli (i.e., linear gratings and phase scrambled). Due to this dissociation of processes acting on a target when visible versus invisible, it would be interesting to investigate them systematically in states of awareness and unawareness.

The asymmetry between entering and exiting awareness was further illustrated in the behaviour of observers’ pupils. Overall, the pupil of the target eye and mask eye paralleled each other (**Figure S1a,b**), exhibiting consensual responses as expected of normal function (Akova et al., 2025). As an expected consequence of target contrast ramping, pupillary contrast responses were observed (Fitzpatrick et al., 2024); pupils constricted as target contrast increased and dilated as contrast decreased (**Figure S1**). However, the contrast-induced dilation plateaued, and constriction of the pupils occurred as targets were approaching their exit from awareness (**Figure 4a,b**), despite the continued decreasing of target contrast. This is similar to the pupillary constriction seen as the offset of static sun images within awareness is approached (Sperandio et al., 2018). In further agreement with Sperandio et al. (2018), observers’ pupils displayed a small initial dilation before a sustained constriction when target stimuli were flashed suppressed from awareness (**Figure S1c,d**). In comparison, observers’ pupils dilated both before and after they reported unawareness of the target (**Figure S2a,b**). Prior to the report, dilation is expected as a pupillary contrast response to the decreasing target contrast. The continued dilation after is not explainable by a simple physiological response. As such, it is more likely that these periods of pupil dilation are in response to the transition from awareness to suppression at the corrected time point (Sperandio et al., 2018).

Pupil constriction prior to a perceptual switch and followed by dilation is also seen in binocular rivalry (Acquafredda et al., 2022). This change did occur around reported breakthrough points (**Figure S2c,d**), but not around corrected breakthrough points (**Figure 4c,d**). However, it is not possible to disambiguate between a contrast response and a genuine transition from target unawareness to awareness as the contrast ramping of the target reversed at reported breakthrough. To separate the processes, a study that stops or does not reverse the contrast ramping at reported breakthough could be conducted. Without the ability to verify, the corrected breakthrough time point is not necessarily invalidated. The onset of detection targets induces pupil dilation that seemingly scales with expected task engagement (Privitera et al., 2010). While we did not observe dilation after corrected breakthrough times, the delay in contrast-induced constriction for the orientation discrimination may be the result of a larger task dependent dilation acting in opposition (**Figure S1d**). A future study could verify this by measuring the pupillary response in a tCFS paradigm without any task or by employing more engaging tasks (e.g., identifying complex features). From these results and those of previous studies, pupil dilation appears to hold promise as a physiological marker of (un)awareness.

Inherent to the contrast ramping in tCFS and bCFS is the “overshoot” of thresholds measured due to the continued contrast change during response times. The current study succeeded in quantifying these response time components and in doing so, revealed the asymmetry in their contributions to breakthrough and suppression thresholds. Taking this asymmetry into account, a mathematical model for correcting thresholds was developed. Changes in pupillary dynamics found for perceptual switching were observed around these corrected time points, supporting their validity. Prior to and after correcting for response times, the contrast increase required for a target to break through from suppression (i.e., suppression depth) was less than the decrease in contrast for the same target to return to suppression (i.e., suppression resistance). Together, these findings open a wide range of future directions for the tCFS paradigm, providing a physiological marker for transitions in awareness and highlighting its ability to investigate phenomena in both states of awareness and unawareness.

## Acknowledgements

This work was supported by an Australian Research Council (ARC; Australia) Discovery Project Grant (DP250100118) awarded to RR and DA and a National Health and Medical Research Council (NHMRC; Australia) Investigator Grant (2026318) awarded to RR.

## Supplementary Materials

**Figure S1.**
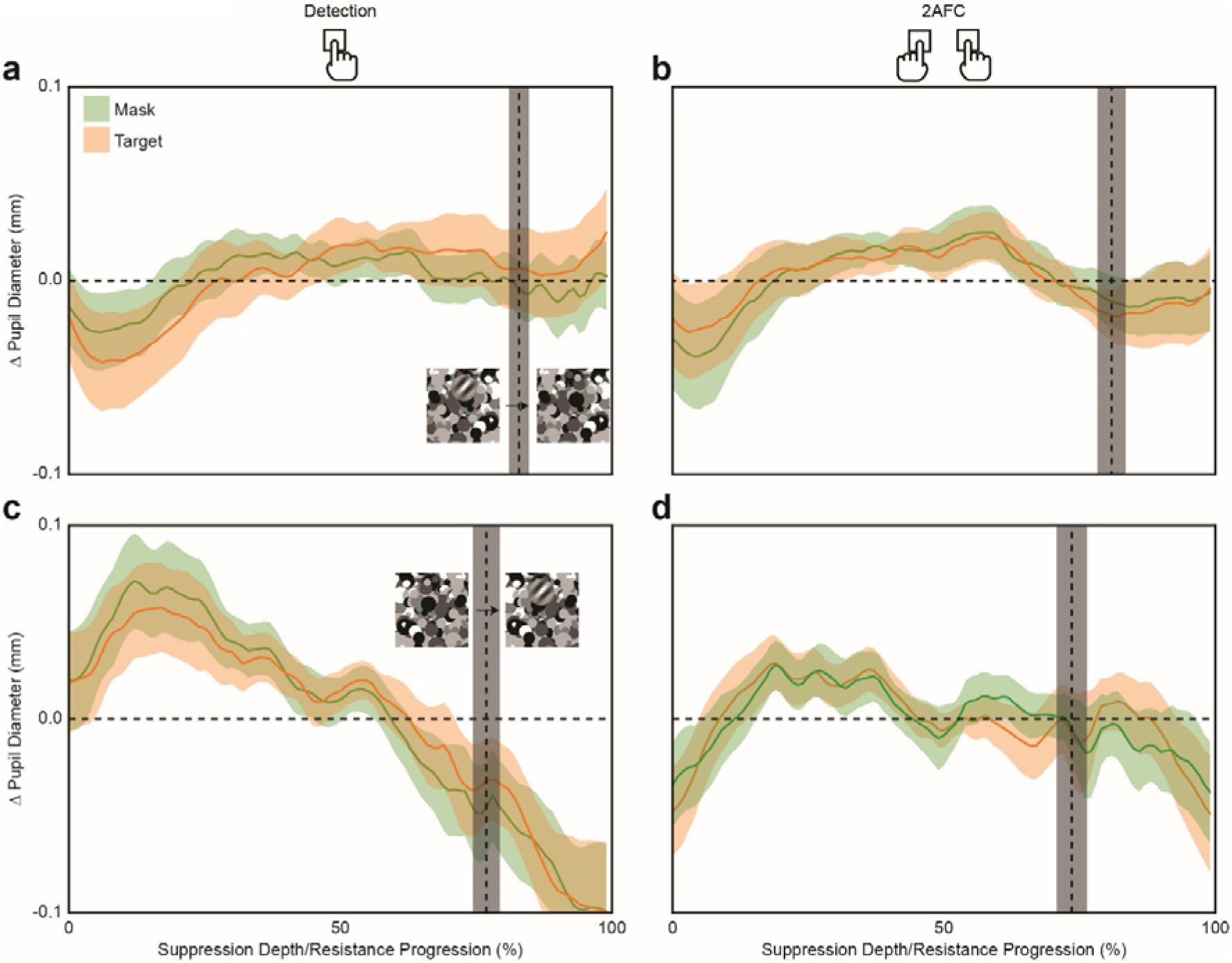
Pupil Diameter as a Function of Normalised Progression Between Breakthrough and Suppression. (**a**) Displays the change in pupil diameter of the mask- (green) and target- (orange) presented eyes as the target entered suppression while the participant was tasked with detecting the target on breakthrough. (**b**) Same as (**a**), but when the participant had reported the orientation of the target when it broke through. (**c-d**) Same as (**a-b**), but when the target was exiting suppression and entering awareness. The time between successive breakthrough and suppression points was normalised and taken as a measure of depth progression. Vertical dashed line in (**a, b**) and (**c, d**) indicates the corrected point of suppression and breakthrough, respectively. Shaded regions indicate ± SEM.

**Figure S2.**
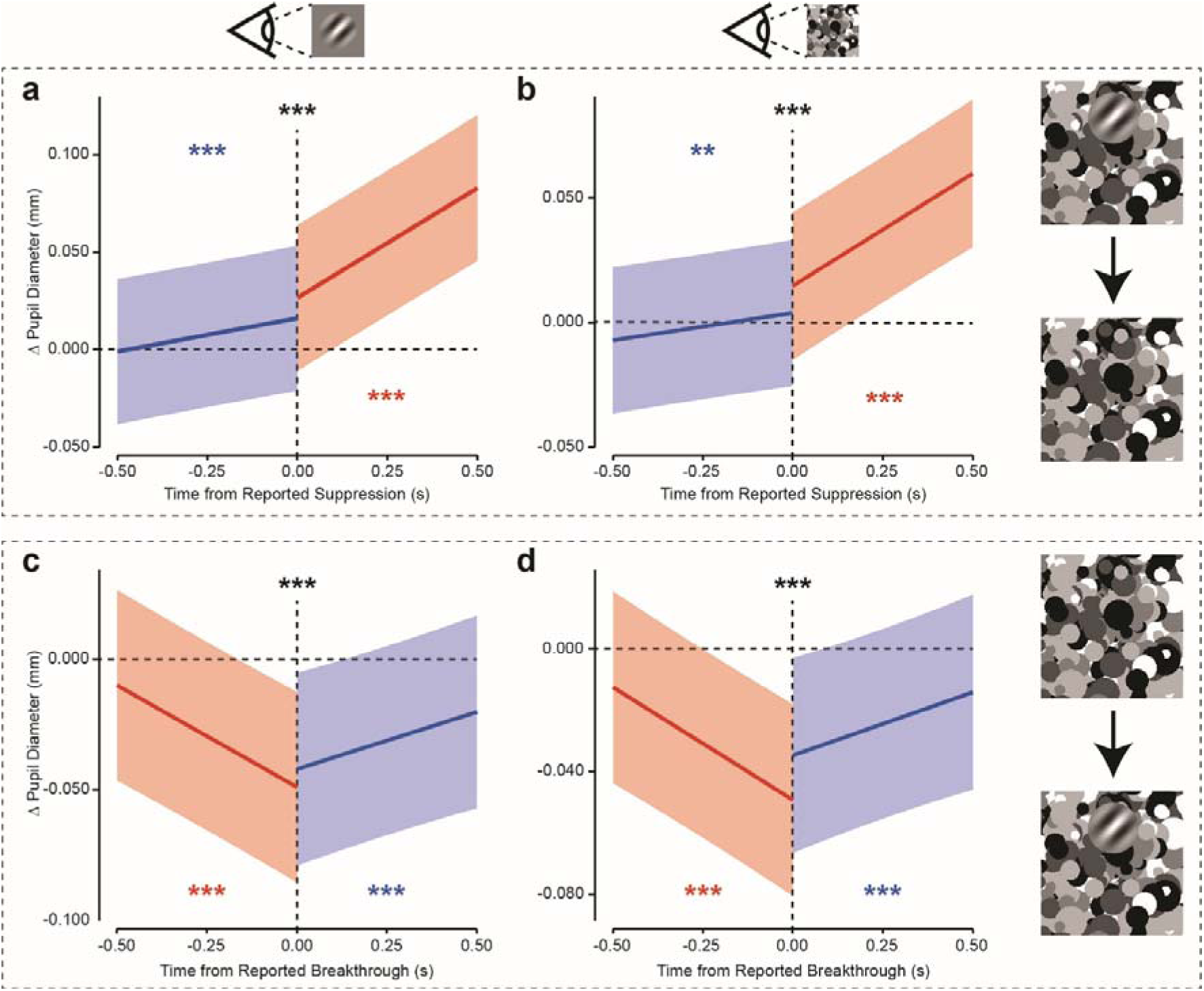
Pupil Diameter as a Function of Time Around Reported Suppression and Breakthrough. (**a**) Displays the estimated fixed effect of time on pupil diameter of the target presented eye 500 ms before and after the reported point of transition from conscious awareness to suppression of targets. (**b**) Same as (**a**), but for the eye presented with the Mondrian mask. (**c-d**) Same as (**a-b**), but for the reported point of transition from suppression to conscious awareness. Coloured lines and regions correspond to the reported state of conscious awareness of the target; blue for targets within awareness and red for suppressed targets. Shaded regions represent 95% confidence intervals. \**p* < .05. \*\**p* < .01. \*\*\**p* < .001, where coloured significance indicators are for corresponding slopes against 0 and black indicators are for differences in slopes across transition point. Shifts between states are indicative of linear fits that do not align with the pupil diameter at time 0 s.

